# pMAT: An Open-Source, Modular Software Suite for the Analysis of Fiber Photometry Calcium Imaging

**DOI:** 10.1101/2020.08.23.263673

**Authors:** Carissa A. Bruno, Chris O’Brien, Svetlana Bryant, Jennifer Mejaes, Carina Pizzano, David J. Estrin, David J. Barker

**Affiliations:** Department of Psychology, Rutgers, The State University of New Jersey; Feil Family Brain & Mind Research Institute, Weill Cornell Medicine

**Keywords:** Fiber Photometry, Calcium Imaging, Neural Recording, Open Source

## Abstract

The combined development of new technologies for neuronal recordings and the development of novel sensors for recording both cellular activity and neurotransmitter binding has ushered in a new era for the field of neuroscience. Among these new technologies is fiber photometry, a technique wherein an implanted fiber optic is used to record signals from genetically encoded fluorescent sensors in bulk tissue. Fiber photometry has been widely adapted due to its cost-effectiveness, ability to examine the activity of neurons with specific anatomical or genetic identities, and the ability to use these highly modular systems to record from one or more sensors or brain sites in both superficial and deep-brain structures. Despite these many benefits, one major hurdle for laboratories adopting this technique is the steep learning curve associated with the analysis of fiber photometry data. This has been further complicated by a lack of standardization in analysis pipelines. In the present communication, we present pMAT, a ‘photometry modular analysis tool’ that allows users to accomplish common analysis routines through the use of a graphical user interface. This tool can be deployed in MATLAB and edited by more advanced users, but is also available as an independently deployable, open-source application.

## 1.0 Introduction

Fiber photometry is a tool that allows for the recording of bulk fluorescent signals from a growing number of sensors including those for detecting levels of cellular activity, such as GCaMP6 (Chen et al., 2013), jGCaMP7 (Dana et al., 2019), jRCAMP1a,b or jRGECO1a (Dana et al., 2016). In addition, a number of sensors have recently been developed for detecting the binding of specific neurotransmitters, such as glutamate via iGluSnFr (Marvin et al., 2018), GABA via iGABASnFr (Marvin et al., 2019), acetylcholine via iAChSnFr (Borden et al., 2020), serotonin via iSeroSnFr (Unger et al., 2019), dopamine via dLight (Patriarchi et al., 2018) or GRAB_DA_ (Sun et al., 2018), and norepinephrine via GRAB_NE_ (Feng et al., 2019).

A major advantage of fiber photometry is its ease of use and capability to conduct high-throughput experiments. Recordings are accomplished via lightweight fiber-optics that provide little interference in most behavioral apparatuses. Moreover, the inclusion of an internal control channel (i.e., an ‘isosbestic control’ channel) (Lerner et al., 2015)) provides a clever method for subtracting movement artifacts that are generated during the task. Fiber photometry can reliably record activity at cell bodies (Barbano et al., 2020) as well as axon terminals (Barker et al., 2017) and has proven especially useful for repeated or longitudinal recordings (Li, Liu, Guo, & Luo, 2019; Pignatelli et al., 2020) or for recording from deep-brain structures (Root & Barker, 2020) Lastly, fiber photometry provides the ability to conduct circuit-specific recordings. This can be accomplished by injecting viral vectors (e.g., GCaMP) at the soma for a target group of cells while placing the fiber optic over distally located axon terminals (Barker et al., 2017) or by injecting retrograde viruses (e.g., retrograde adeno-associates or herpes simplex viruses) at the site of axon terminals and placing the fiber optic over the cell bodies from which those terminals originate (Barker et al. unpublished data).

Following a landmark paper by Gunaydin and colleagues (2014), there has been an exponential growth in the use of fiber photometry within the neuroscience field (Figure 1). Accordingly, there has been a concomitant increase in the number of open-source (e.g., Akam & Walton, 2019) or commercially available systems for the acquisition of fiber photometry data, including recent movements towards wireless systems. Nonetheless, the analysis of fiber photometry data is most often accomplished via custom programming scripts, creating a barrier for many laboratories as they struggle to tackle the analysis of the large datasets that can be amassed using the technique. Additionally, efforts to standardize methods of analysis for fiber photometry have, to date, mostly existed at the grassroots level, while multiple analysis packages already exist for single-cell or two-photon calcium imaging. Thus, the goal of the present manuscript was to develop an open-source analysis suite capable accomplishing basic tasks required for the visualization and analysis of fiber-photometry data. The present Photometry Modular Analysis Tool or ‘pMAT’ can be expanded upon via future MATLAB programming, but has also be deployed as an executable file (‘.exe’) that runs independent of MATLAB. In addition to alleviating the programming barrier to entry, the analysis suite provides a starting point for the standardization of analysis protocols and can be expanded to support analysis for most of the common data acquisition systems currently on the market.

**Figure 1.**
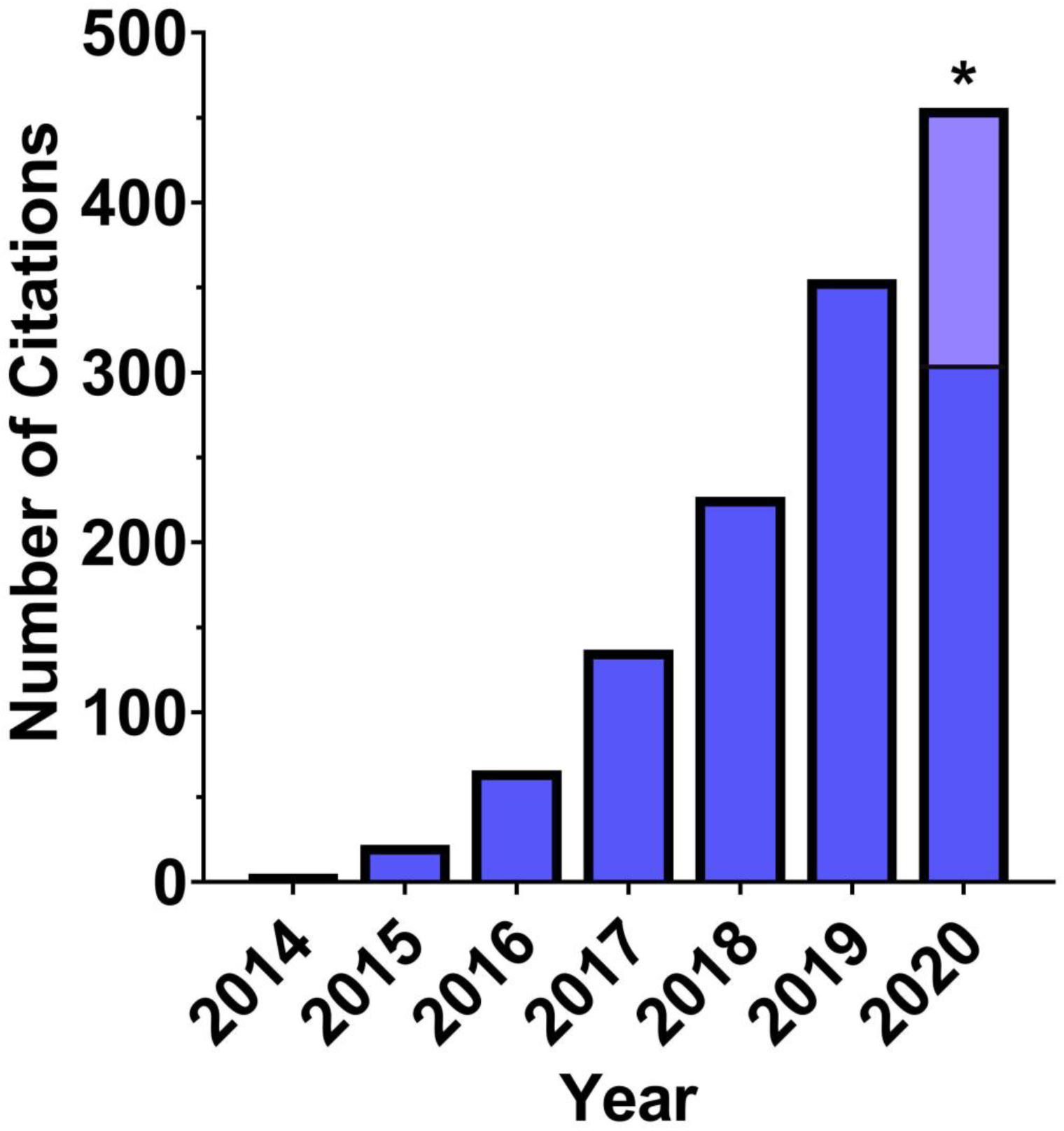
Growth in the use of fiber photometry calcium imaging since its inception. The implementation of fiber photometry has increased rapidly since 2014 and continues to grow yearly at a rapid pace. The data here represents the number of Google Scholar references to Fiber Photometry. * Projected 2020 papers, light blue; current, dark blue.

## 2.0 Recording Parameters for Testing

For all internal recordings using to validate the software pipeline, GCaMP6 was excited at two wavelengths, a ∼490nm, calcium-dependent signal and ∼405 nm isosbestic control (Lerner et al., 2015), by amplitude modulated signals from two light-emitting diodes reflected off dichroic mirrors and coupled into a 400µm 0.48NA optic fiber. The isosbestic control channel (henceforth termed, ‘control’) represents the point of GCaMP excitation at which emissions from the calcium bound and unbound state have equal intensities. Thus, the control channel represents GCaMP emissions that are independent of calcium, but still susceptible to artifacts related to movement, fiber bending, and all other physical alterations. Signals emitted from GCaMP6s and its isosbestic control channel then returned through the same optic fiber and were acquired using a femtowatt photoreceiver (Model 2151; Newport), digitized at 1kHz, and then recorded by a real-time signal processor (RZ5D) from Tucker Davis Technologies (TDT) running the Synapse software suite. Behavioral inputs were collected using the digital input and output ports (DIO) using either a DIG-726TTL-G card (Med-Associates Inc.) that was connected to the TDT system with a custom-soldered DB-25 cable or using an SG-231 28V DC to TTL adapter (Med-Associates Inc) connected to the TDT system using a BNC connection. Both setups has been tested for temporal precision to verify the synchrony of behavioral data and fiber photometry data. The temporal prevision was always of <1 ms (∼250 µs)

## 3.0 Installation of the pMAT suite

### 3.1 Installation of pMAT suite in MATLAB

For installation in MATLAB, pMAT should be downloaded from https://github.com/djamesbarker/pMAT and saved to a Windows PC. It is recommended that users create a directory called “*C:\pmat-BarkerLab\*”, as this is the location that will be used for saving pMAT settings files. Once downloaded, it is important to open MATLAB and use the *pathtool* command (type ‘*pathtool’* into the command window) to add and save the pMAT folder and all of its subfolders to the MATLAB path. pMAT is then run by typing *pmat* into the MATLAB command window. pMAT was developed with MATLAB 2019a, so we recommend using this version or newer.

### 3.2 Installation of pMAT as a deployable .exe file

To run pMAT as an independent ‘.exe’ file, one of the pMAT ‘independent deployment’ folders should be downloaded from https://github.com/djamesbarker/pMAT. The folder should be unzipped and saved to your computer. Open the pMAT deployment folder and click the application named ‘pmat installer’. The default path is ‘*C:\Program Files\pmat-BarkerLab*’. This process will install both the pMAT software and the MATLAB runtime environment, which allows you to use the program as an executable program (*‘*.*exe’*) without the need for a MATLAB license. At the time of this manuscript, the MATLAB 2020a runtime (i.e., runtime 9.8) is being used for deployment.

## 4.0 Features of the pMAT suite

### 4.1 Data Loading and Control Module

The pMAT system was initially designed around the Tucker Davis Technologies (TDT) recording platform, as the original fiber photometry systems were run on this hardware and software, which provides for synchrony between behavioral data and neural signals. Calcium or neurotransmitter based signals recorded from these experiments are stored in data ‘tanks’ and ‘blocks’. tanks often hold groups of recordings (e.g., multiple recordings from one experiment or subject), while the blocks contain each individual recording session. The first step in loading data is to import data. pMAT allows you to process either single files or whole folders full of files using ‘batch processing’. If selecting a single file, you will be prompted to select the specific TDT block containing your data. If selecting batch processing, an entire folder (containing many TDT data tanks) can be selected for processing. In addition, data from other systems can be loaded into the pMAT suite via the ‘Import Data’ menu. Users import two CSV files, one containing the signal and control channel, as well as their relevant timestamps, and the other containing unique strings identifying different behavioral events, as well as their onset and offset times. This format accommodates many other recording systems on the market, although additional import tools are also planned in future releases. Upon loading a file, the user will be prompted to buffer the file by cutting out the first *x* seconds. This buffer is meant to account for the rise or fall of the light-emitting diodes (LEDs) used in the system. The default is set to 2000 samples (∼2 seconds in most systems). Once a file has been loaded, users can append additional even data (e.g., video scored events) from a ‘.*csv’* file using the ‘Import Data’ menu. Additionally, loading a single file allows the user to interact with the other modules in the pMAT suite. Each module can be run independently using buttons within each module, or the entire suite can be set up at once and executed using the ‘Run all Selections’ button. A similar approach is taken for batch processing. The user interacts with the first file to set up the parameters for batch processing and then runs all of the files in the folder by pressing the ‘Run All Selections’ button (Figure 2A).

**Figure 2.**
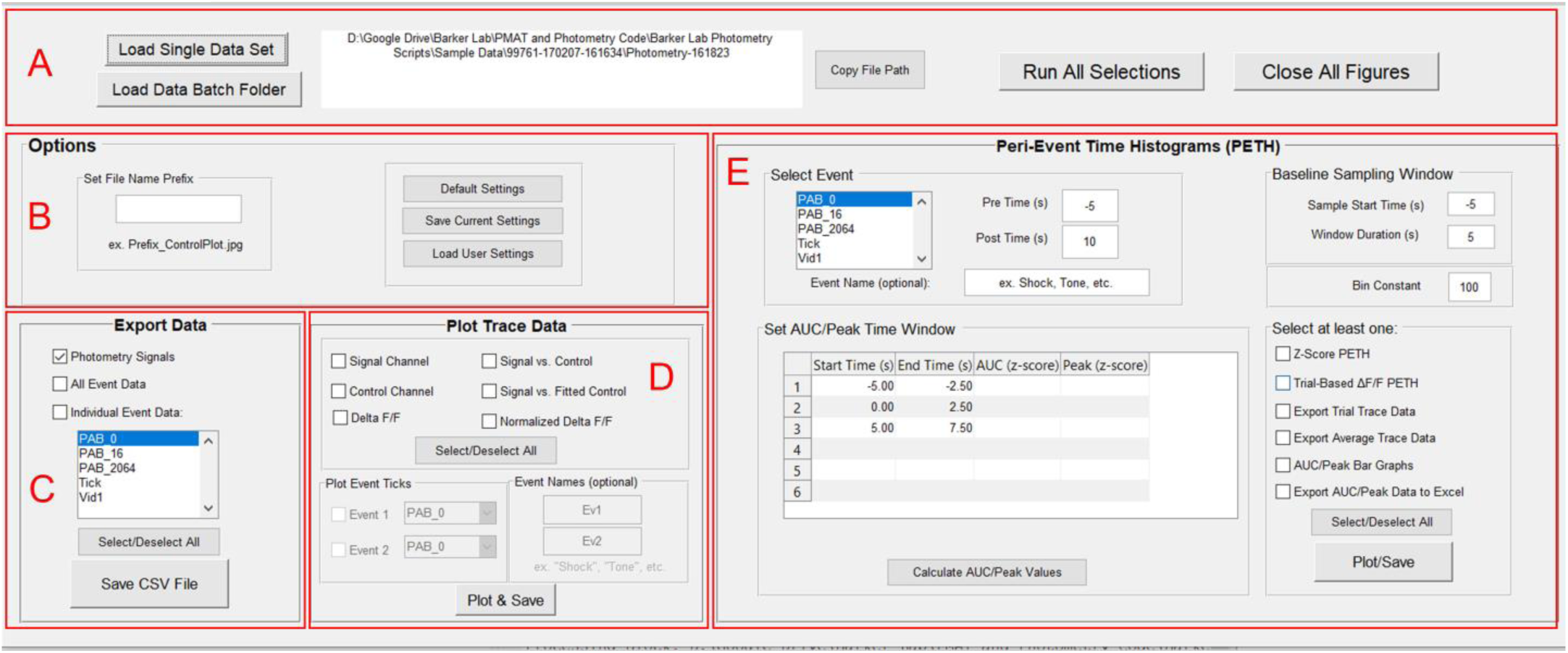
Overview of the pMAT suite. **A)** The control module allows for loading single files or batch processing and provides an interface of buttons to run selections across all modules or to close all of the figures generated when running the code. **B)** The Options module is used for setting file prefixes as well as saving and loading pMAT options. **C)** The Export Data module provides options for exporting signal, control, or event data into comma separated value (‘.*csv’*) files that can be imported into other programs such as Microsoft Excel or R. **D)** The Plot Trace Data module allows for the plotting of signal and control channels, as well as the stepwise evaluation of the signal correction routines used to generate the ΔF/F. The module also allows for the visualization of events across time on the ΔF/F plots. **E)** The Peri-event time histogram module allows for the evaluation of the signal around an event of interest and provides the tools for the plotting and exporting of relevant data for the peri-event traces or summary statistics.

### 4.2 Options Module

Settings for the state of the pMAT suite can be saved by the user for future use, or for collaboration across laboratories (Figure 2B). These files, by default, will be saved in the “*C:\pmat-BarkerLab\”* folder, unless directed elsewhere, and can be loaded again at a later date. Settings that can be saved include file prefixes, checkbox statuses, list box selections, popup selections, as well as all user input text boxes (pre/post times, baseline sampling window and bin constant). The user can also set a file prefix to help organize data files (e.g., by adding an experiment name, dosing group, etc.) or to help differentiate files (e.g., by adding a version number). Upon closing, pMAT will automatically prompt users by asking whether they wish to save these settings (Figure 3A-D).

**Figure 3.**
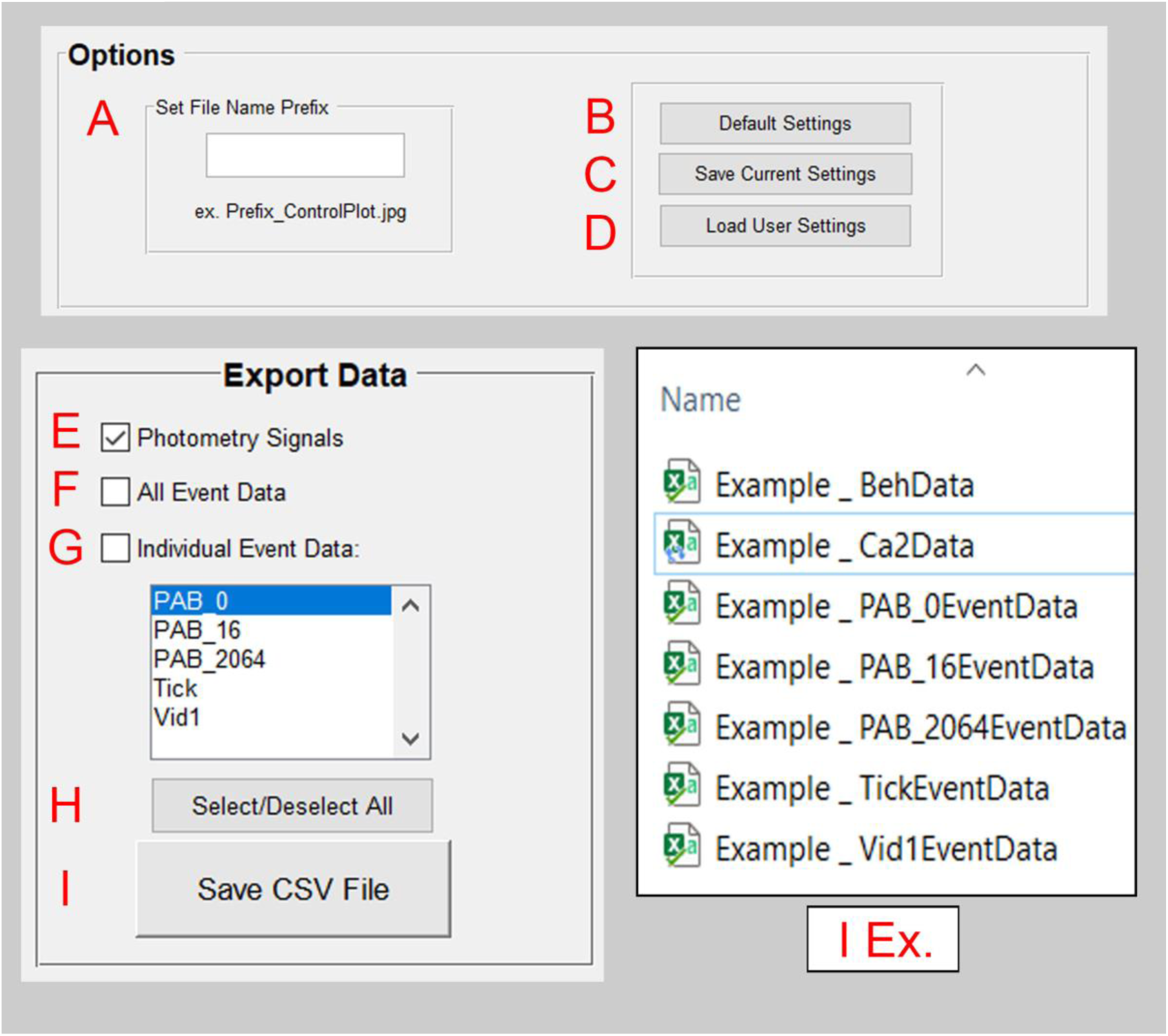
The pMAT options and Export Modules. **A)** Within the options module, a file prefix can be set to help with the organization of files. **B)** The ‘Default Settings’ button reverts all settings back to their original state. **C)** The ‘Save Current Settings’ button allows for users to save settings for the state of the pMAT suite. Users will also be prompted to save their settings when exiting the program. **D)** The ‘Load User Settings’ button allows users to restore previously saved settings, allowing for the reproducibility of routines. **E)** The ‘Photometry Signals’ check box will store a *‘*.*csv’* file with the signal and control channel data as well as their corresponding timestamps, once the ‘Save CSV File’ button (I) is pressed. **F)** The ‘All Event Data’ check box will store a *‘*.*csv’* file containing the tags for all behavioral events as well as their corresponding timestamps, once the ‘Save CSV File’ button (I) is pressed. **G)** The ‘Individual Event Data’ check box will store a *‘*.*csv’* file for each event that has been selected within the window below, once the ‘Save CSV File’ button (I) is pressed. Multiple events can be highlighted for export simultaneously. **H)** Users can select or deselect all checkboxes simultaneously. **I)** The ‘Save CSV File’ button executes the command to save data corresponding to the selected checkboxes in E-G.

### 4.3 Data Export Module

At the most basic level, the pMAT suite allows the user to quickly export data in a universal ‘.*csv’* format that is compatible with Microsoft Excel, MATLAB, and Python, among others (Figure 2C). Three types of data can be exported: 1) a file containing calcium signals, control signals, and their corresponding timestamps, 2) a file containing all event data and timestamps, which correspond to all transistor-transistor logic (TTL) inputs to the recording hardware for behavioral or experimental events, and 3) selected event data and their corresponding timestamps. Each of these files are saved in the “Data” folder created within the corresponding data block. Currently, the storage location of data is predetermined in order to facilitate the batch processing of data (Figure 3E-I).

### 4.4 Plot Trace Data Module

Traces across whole experimental sessions are constructed using a multi-step process. Each of these steps can each be individually visualized and evaluated by users with the tools provided in the ‘Plot Trace Data’ module of the GUI (Figure 2D). First, data from the signal and isosbestic control channels are extracted. Second, the channels are individually smoothed using a Lowess, local linear regression method to smooth the signal and reduce high-frequency noise (signal channel and control channel plots; Figure 4 A-C). As signal and control channel emissions generally have different levels of power (‘Signal vs. Control’ plot), the next step is to normalize the scale of the channels. This is accomplished by first using a least-squares regression to find the relationship between the signal and control channels (MATLAB *polyfit* function):

**Figure 4.**
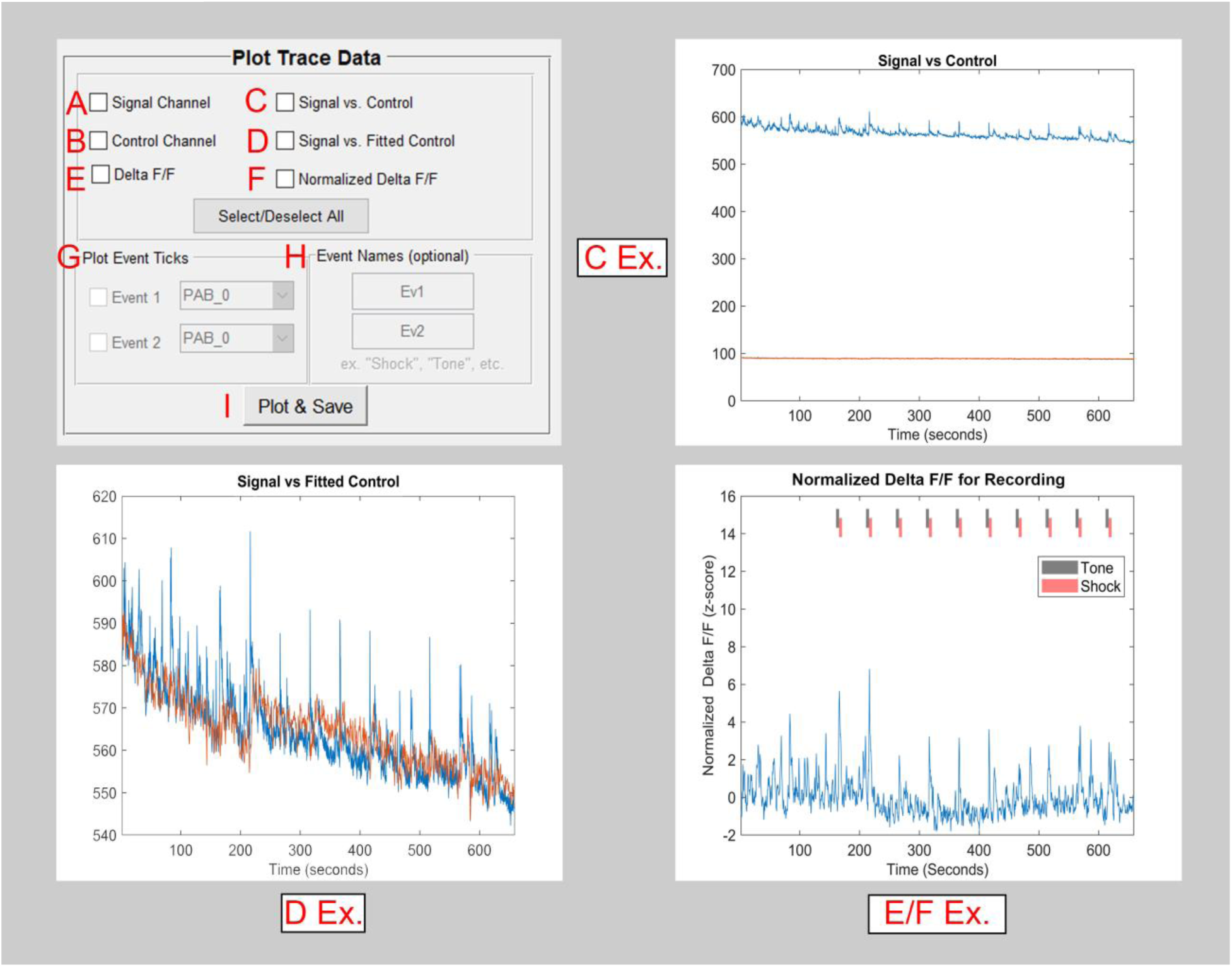
The pMAT trace plotting module. **A)** The checkbox for the ‘Signal Channel’ will independently plot data from a recorded sensor. **B)** The checkbox for the ‘Control Channel’ will independently plot data from an isosbestic or autofluorescence control. **C)** The signal and control channels can be plotted simultaneously in order to compare the various properties of these signals (e.g., rate of photobleaching). **D)** The fitted control is overlaid onto the signal channel (D Ex.) in order to help visualize the scaling that has occurred and the corrections that will be made. **E)** The Δ F/F is generated using the signal and fitted control. This shows the final stage of the calculated correction that occurs using the control channel. **F)** A normalized (z-score) version of the Δ F/F is generated. **G)** The event checkboxes allow for the visualization of events across the whole-session trace data. (e.g., the Tone and Shock ticks in the E/F Ex.) **H)** Events can be given a custom name, as desired. **I)** All checkboxes selected in the Plot and Trace Data module are run using the Plot and Save button.

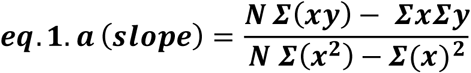

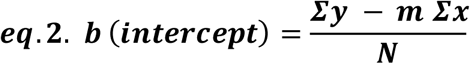

Where N is the number of subjects, x is the control channel, and y is the signal channel. Next, the resulting slope and intercept are then used to generate a scaled control channel (‘Signal vs. Fitted Control’ plot; Figure 4D):

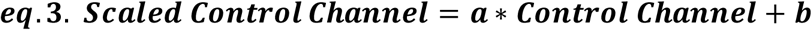

Finally, the delta F/F (ΔF/F) is generated by subtracting the fitted control channel from the signal channel. This subtraction is used to eliminate movement or other common artifacts (Figure 4E):

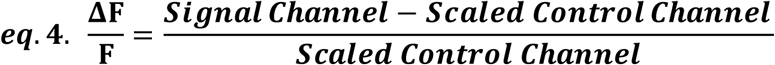

For the fully processed ΔF/F plots, the user also has the option to represent the whole session trace as a normalized z-score (z-score of eq. 4) of the trace and to include reference ticks showing the timing of up to two behavioral events on top of the whole-session trace (Figure 4 G-H).

### 4.5 Peri-Event Histograms Module: Overview

Peri-event time histograms/heatmaps (PETHs) are a valuable tool for examining the stability of a response over repeated trials. To calculate PETHs in the pMAT suite, a user-defined event window is set around each event that begins ***x*** seconds before the event and ends ***y*** seconds after the event. In addition, a baseline window is defined for a user-defined period of time preceding each event window. For both the baseline window and event window, the ΔF/F is then calculated in the same manner as described above for the trace data (Section 4.4), albeit on a much shorter timescale. In addition, the data are centered following the ΔF/F calculation to set the first point in each trial to zero. These data are visualized using the “Trial-Based ΔF/F PETH” feature.

The combination of this shortened correction window and zeroing procedure are designed to correct for changes in the stability of the signal that occur over long timescales due to photobleaching of fluorophores, photobleaching of fiber optic cables, and changes in the stability of other components of all photometric systems. The challenge of a moving signal is not unique to fiber photometry and has been effectively solved in electrochemistry experiments for some time using a similar approach (Rodeberg et al., 2017). Signals are then converted into a robust z-score using the following formula:

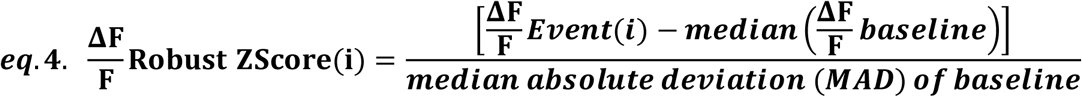

Traces for both the average trace, or the traces from individual trials can then be exported for further analysis or for plotting elsewhere.

### 4.6 Peri-Event Histograms Module: Area under the curve and peak calculations

From the robust z-score, the area under the curve and peak values can be extracted from the PETH data for up to six different user-defined windows. These windows are defined by editing the cells in the “Set AUC/Peak Time Window” matrix.

The peak is calculated as the maximum value that occurs within each user-defined window. The area under the curve is calculated for the portion of the curve that falls within each user defined window using the trapezoidal method for integral calculation (MATLAB trapz function). These data can be plotted as a bar graph for examination and also exported in a ‘.csv’ format for further analysis.

### 4.7 Sensor Compatibility

We are currently in an era where novel sensors are being developed at a rapid pace. Many but not all of these sensors have been tested using fiber-photometry in their original publications. With this in mind, we sought the help of numerous labs in order to provide an up-to-date list of the tested and un-tested sensors currently being used with fiber photometry. We also used this opportunity to evaluate these sensors within the pMAT suite, to ensure its broad compatibility. The list of sensors and relevant publications are found in Table 1, with plots in Figure 6.

**Table 1.** Genetically Encoded Sensors. A list of genetically encoded fluorescent sensors, including the most up-to date information regarding whether these have been used effectively for fiber-photometry calcium imaging.

| Sensor Name | Validated for<br>Fiber<br>Photometry | Relevant Citations | Lab/Individual<br>contributing validation<br>data or validating citation |
| --- | --- | --- | --- |
| GCaMP6 (f,m,s) | Y | Gunaydin et al., 2014 | Barker Lab (David Barker) |
| jGCaMP7 (b,s,f,c) | Y | Dana et al., 2019 | Rothwell Lab (Marc Pisansky) |
| jRCaMP1a,b or jRGECO1a | Y | Dana et al., 2016; Kim et al., 2016 | Lerner Lab (Jillian Seiler) |
| dLight-1.1 | Y | Patriarchi et al., 2018 | Lerner Lab (Jillian Seiler); Papaleo Lab (Daniel Dautan) |
| iGluSnFR | Y | Marvin et al., 2018; McGovern et al., 2020 | Root Lab (Dillon McGovern) |
| iGABASnFR | Y | Marvin et al., 2019; McGovern et al., 2020 | Root Lab (Dillon McGovern) |
| iAChSnFR | Y | Borden et al., 2020; McGovern et al., 2020 | Root Lab (Dillon McGovern) |
| iSeroSnFR | TBD | Unger et al., 2019 | Lin Tian (Chunyang Dong) |
| GRAB<sub>DA</sub> | Y | Sun et al., 2018 | Cheer Lab (Kate Peters) |
| GRAB<sub>NE</sub> | Y | Feng et al., 2019 | Feng et al., 2019 |
| GRAB-5HT | Y | Wan et al., 2020 | Wan et al., 2020 |
| iATPSnFR | TBD | Lobas et al., 2019 |  |
| iGlucoSnFR | TBD | Keller et al. 2019 |  |

**Figure 5.**
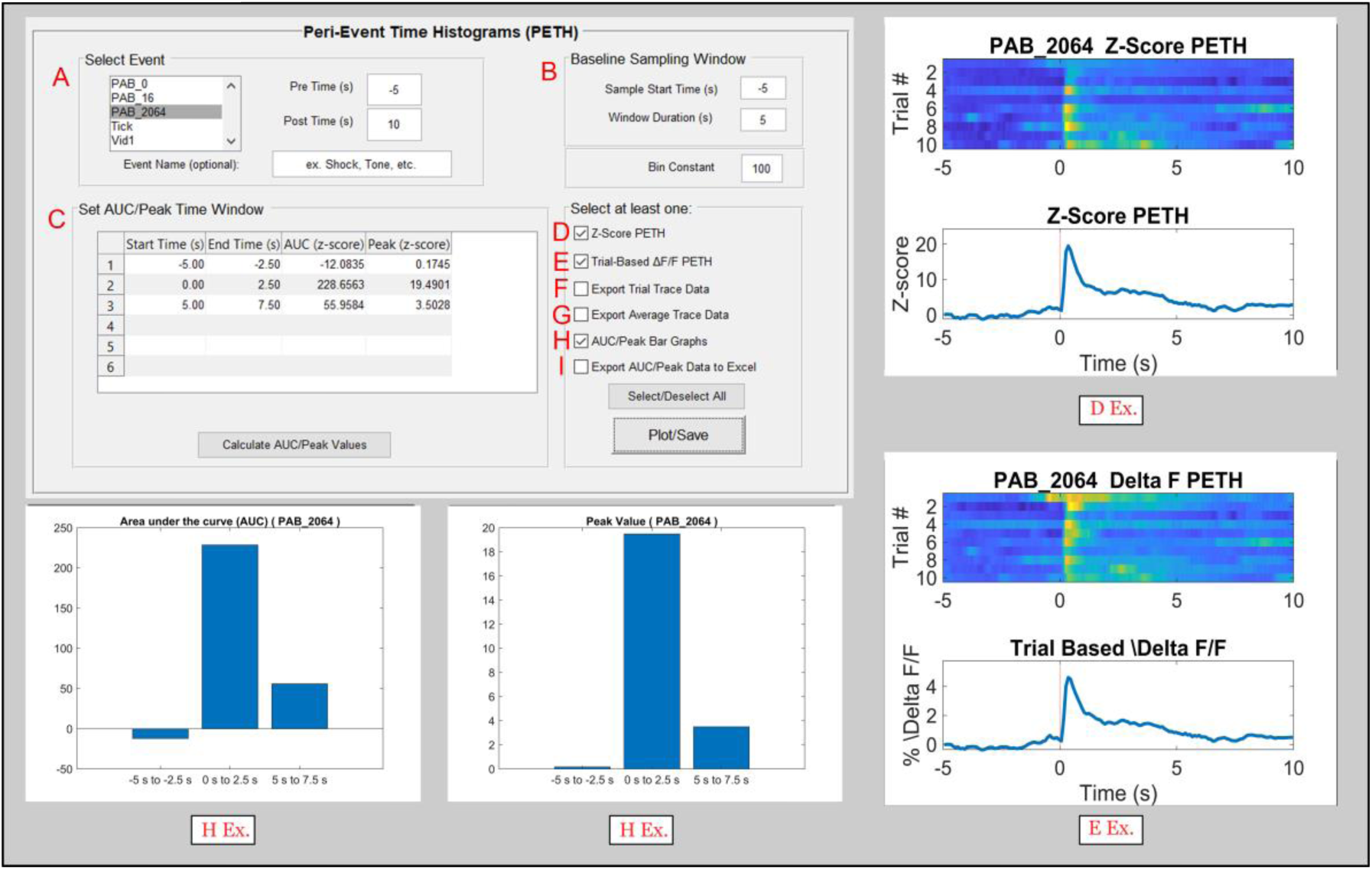
The pMAT peri-event time histogram module. **A)** An event of interest can be selected to create a peri-event heatmap and histogram (PETH; D Ex. And E Ex.). A window of event before (Pre Time) and after (Post Time) can also be defined. **B)** The user can then define a Baseline Sampling Window to act as a reference point for normalizing (robust z-score) data. In addition, a bin size for the event and baseline data can be defined. **C)** The area under the curve (AUC) and ‘Peak’ or maximum value of a trace are two commonly used metrics collected for fiber photometry data. Users can input up to 6 values for the start and end times of specific windows and the AUC and Peak will be calculated for each of these windows. These values can also be plotted using the checkbox in H. **D)** Selecting the ‘Z-Score PETH’ (peri event histogram and heatmap) will plot a normalized (robust z-score) heatmap and histogram aligned to the event selected in A. Warmer colors on the heatmap (top) represent higher fluorescent signals at that timepoint, while the line plot on the bottom represents the average for all trials corresponding to the event of interest (D Ex.). **E)** The ‘Trial Based Δ F/F PETH’ will plot the absolute Δ F/F values on a peri event heatmap and histogram (E Ex.). **F)** Data for individual trials of the Z-Score PETH (D; top) can be exported to a ‘.*csv’* file and saved for plotting or processing elsewhere. **G)** Data for the average trace of the Z-Score PETH (D; bottom) can also be exported to a *‘*.*csv’* file for plotting or processing elsewhere. **H)** The checkbox for H produces two graphs, showing the AUC and Peak values defined by the user in the cells provided in C. **I)** The AUC and Peak values can also be exported into a ‘.*csv’* file. These are commonly collated across experimental subjects or groups for statistical analysis.

**Figure 6:**
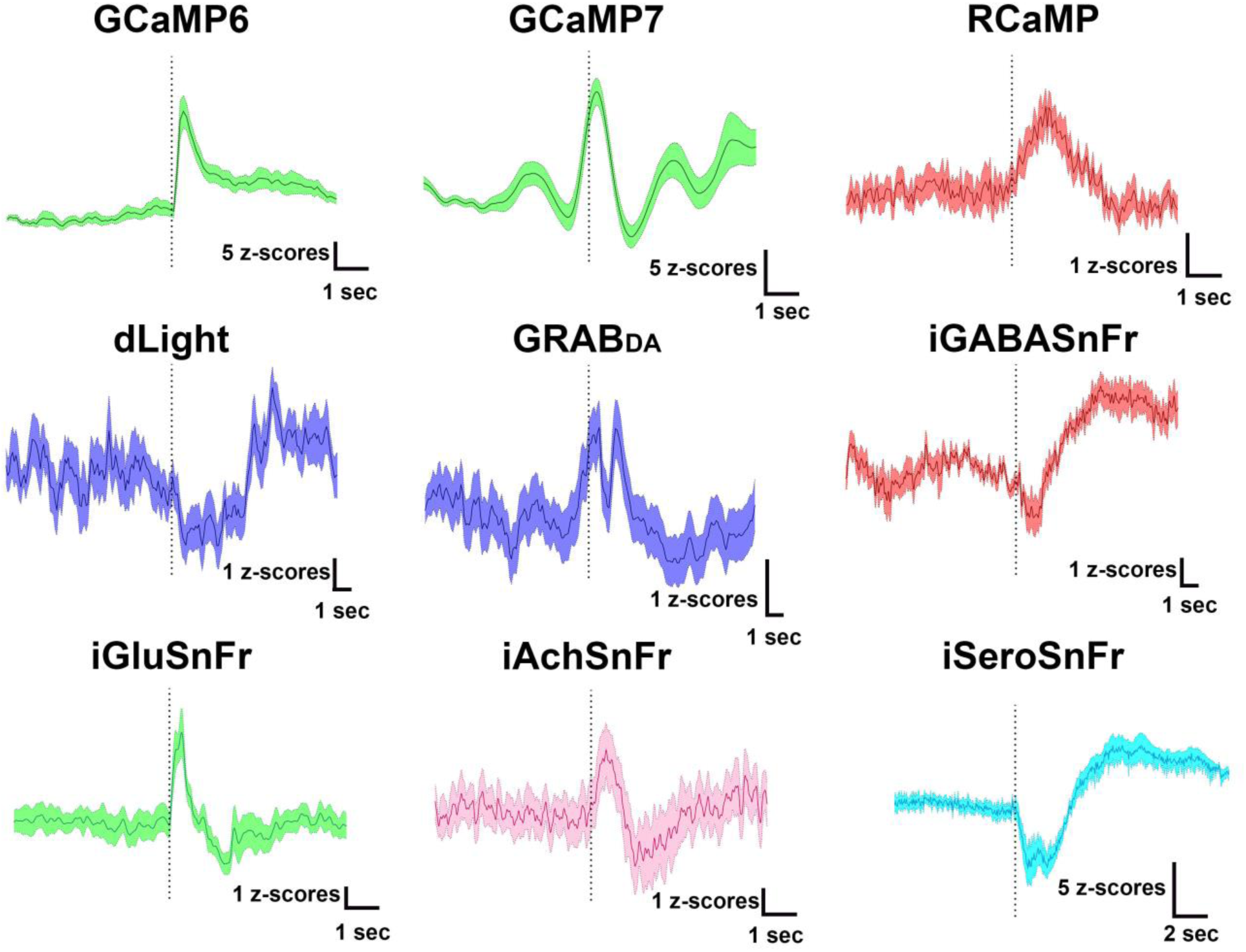
Examples of sensors compatible with fiber photometry calcium imaging. Example traces from sensors used to detect cellular activity (GCaMP6, GCaMP7, or RCaMP) or to detect neurotransmitter binding for dopamine (dLight, GRAB_DA_), GABA (iGABASnFr), glutamate (iGluSnFr), acetylcholine (iAchSnFr), or serotonin (iSEroSnFr). The events represented are different for each sensor, but all are aligned to a relevant experimental event at the point of the dotted line.

### 4.8 Future development goals

The first iteration of pMAT is meant to support the most basic processing with the goal of getting the first iteration of the tool to end-users. However, the second iteration of the pMAT suite is already underway. The current plans for this iteration include a) increased compatibility for additional recordings or data formats, b) the ability to append additional behavioral data from manual scoring or other hardware (e.g., DeepLabCut and SimBA) c) the ability to detrend bleached data without using regression modeling techniques d) the ability to perform transient detection analyses for the quantification of peaks in the signal across entire sessions, e) the ability to generate multi-event histograms for events with fixed or variable intervals, and f) increased compatibility with tools from the TDT Synapse software, such as behavioral notes.

## 5.0 Discussion

Fiber photometry is among a series of recent technological advances that has allowed us to examine levels of neuronal activity, neurotransmitter signaling, and even the interplay between the two that were previously inaccessible in awake, freely moving animals. The ability to conduct fiber photometry recordings has been the product of advances in recording technologies (Gunaydin et al., 2014) as well as advances in fluorescent proteins, which have enabled the development of an entire library of sensors in colors that exist both within and outside of the visible spectrum (O’Banion & Yasuda, 2020).

There are many advantages to the implementation of fiber photometry, including the ability to examine the activity of neurons with specific anatomical and/or genetic identities (Barbano et al., 2020; Barker et al., 2017; Calipari et al., 2016; Fenno et al., 2020; Pignatelli et al., 2020; Root et al., 2020) and the ability to record from multiple cell types or brain regions simultaneously (Dana et al., 2016; Kim et al., 2016; Meng et al., 2018; Sych, Chernysheva, Sumanovski, & Helmchen, 2019). Fiber photometry systems are also relatively simple when compared to other imagine techniques (Siciliano & Tye, 2019), allowing for high-throughput experiments in a wide array of behavioral tasks(Barbano et al., 2020; Barker et al., 2017; Calipari et al., 2016; Pignatelli et al., 2020; Root et al., 2020; Zhong, Li, Feng, & Luo, 2017). Finally, the availability of both open source (Akam & Walton, 2019) and turn-key systems, as well as the relatively low cost of these systems, has made the technique incredibly accessible for most laboratories.

The advantages of fiber photometry must be considered carefully during experimental design and weighed against a number of notable disadvantages. Indeed, fiber photometry captures a relatively homogeneous picture of neuronal activity, which appears biased towards increases in neuronal activity when compared electrophysiological recordings (London et al., 2018), although decreases can be observed when cellular activity is homogeneous (e.g., Zhong et al., 2017; Figure 6). Moreover, fiber photometry carries poor spatial resolution when compared single-cell calcium recordings via miniaturized microscopes or two-photon microscopes (Siciliano & Tye, 2019). Finally, fiber photometry recordings collect data from a fairly small field of tissue (Pisanello et al., 2019), as the collection of these signals is subject to both the dispersion light from low-powered LEDs and the emission of relatively weak signals from calcium and neurotransmitter sensors. Thus, the quality of recordings can be highly dependent on both the proximity of the implant and the density of neurons. This becomes especially important when recording from axon terminals, where the quality of signals are routinely lower than when recording from cell bodies (Barker et al., 2017; Kim et al., 2016).

Perhaps one of the largest barriers for conducting fiber photometry experiments has been the intensive analysis required to obtain data for statistical comparisons. This single barrier complicates the process for researchers that are new to the technique and can make fiber photometry less approachable or even create problems with the rigor and reproducibility of the data. While multiple laboratories and commercial companies have offered guidance or analysis scripts for fiber photometry, the pMAT suite provides the first tool for analyzing fiber photometry data that does not require basic programming skills. Thus, pMAT is a first step towards alleviating this burden and providing an intuitive and accessible way to analyze fiber-photometry data. In addition, having an open-source tool with a well-documented description of the analytical approaches is a first step towards standardizing analysis pipelines. Moreover, the independent deployment of pMAT obviates the need for additional software, making it more widely accessible and budget-friendly. Among the tools available are those to export data for dissemination or plotting, to rapidly check the quality of signals following experimental sessions, and to evaluate the quality of movement controls in a stepwise manner. Moreover, we provide experimenters with the tools to plot sensor responses that are time-locked to behavioral events and to collect summary data that can be used for statistical analysis. Finally, the modular design of this tool allows for the rapid integration of additional tools to accommodate sensor-specific analyses or advances in the analytical approaches used by the field.

## Acknowledgements

We thank Drs. Myles Billard from Tucker Davis Technologies for his critical feedback on the pMAT tool. We also thank Dr. David Root, Dillon McGovern, Dr. Kate Peters, Dr. Joseph Cheer, Jillian Seiler, Dr. Talia Lerner, Dr. Daniel Dautan, Chunyang Dong and Dr. Lin Tian for providing sample data for validation of the various sensors presented in the present manuscript. This work was supported by a NIDA K99/R00 Pathway to independence award (DA043572) and by pilot grants from the New Jersey Alliance for Clinical and Translational Sciences as well as the Rutgers Brain Health Institute. Funding sources were not involved in study design, data collection, and interpretation, or in the decision to submit this research for publication.

## Authors Contributions

DJB conceived this project. DJB, CB, CP, and DJE developed the MATLAB scripts used for PMAT. COB, SB, and JM provided debugging support and helped with figure preparation. DJB, COB, SB and JM wrote the manuscript with the contribution of all authors.

